# Unveiling APOL1 Haplotypes: A Novel Classification Through Probe-Independent Quantitative Real-Time PCR

**DOI:** 10.1101/2023.10.16.562539

**Authors:** Murat Dogan, Christine Watkins, Holly Ingram, Nicholas Moore, Grace M. Rucker, Elizabeth G. Gower, James D. Eason, Anshul Bhalla, Manish Talwar, Nosratollah Nezakatgoo, Corey Eymard, Ryan Helmick, Jason Vanatta, Amandeep Bajwa, Canan Kuscu, Cem Kuscu

## Abstract

**Introduction:** Apolipoprotein-L1 (APOL1) is a primate-specific protein component of high-density lipoprotein (HDL). Two variants of APOL1 (G1 and G2), provide resistance to parasitic infections in African Americans but are also implicated in kidney-related diseases and transplant outcomes in recipients. This study aims to identify these risk variants using a novel probe-independent quantitative real-time PCR method in a high African American recipient cohort. Additionally, it aims to develop a new stratification approach based on haplotype-centric model.

**Methods:** Genomic DNA was extracted from recipient PBMCs using SDS lysis buffer and proteinase K. Quantitative PCR assay with modified forward primers and a common reverse primer enabled us to identify single nucleotide polymorphisms (SNPs) and the 6-bp deletion quantitatively. Additionally, we used sanger sequencing to verify our QPCR findings.

**Results:** Our novel probe-independent qPCR effectively distinguished homozygous wild-type, heterozygous SNPs/deletion, and homozygous SNPs/deletion, with at least 4-fold differences. High prevalence of APOL1 variants was observed (18% two-risk alleles, 34% one-risk allele) in our recipient cohort. Intriguingly, up to 12-month follow-up revealed no significant impact of recipient APOL1 variants on transplant outcomes. Ongoing research will encompass more time points and a larger patient cohort, allowing a comprehensive evaluation of G1/G2 variant subgroups categorized by new haplotype scores, enriching our understanding.

**Conclusions:** Our cost-effective and rapid qPCR technique facilitates APOL1 genotyping within hours. Prospective and retrospective studies will enable comparisons with long-term allograft rejection, potentially predicting early/late-stage transplant outcomes based on haplotype evaluation in this diverse group of kidney transplant recipients.

## INTRODUCTION

Apolipoprotein L1 (APOL1) is a primate-specific apolipoprotein-L family member and is considered as minor component of high-density lipoprotein (HDL). It plays a role in lipid exchange and transport of cholesterol from peripheral cells to the liver. APOL1 gene is on the q-arm of human chromosome-22, comprises five operational domain, along with other five APOL genes (2-6), and encodes several different transcript variants ^1^. APOL1 is primarily synthesized in the liver and found in several tissues such as liver, heart, lung, podocytes and proximal tubules in kidney ^2, 3^.

APOL1 is recognized as a secreted protein that travels through the bloodstream and assembles into a complex referred to as a trypanosome lytic factor (TLF) -which is synthesized from serum resistance-associated binding domain (SRA)^4^-with high-density lipoprotein 3 (HDL3) and the hemoglobin-binding, haptoglobin-related protein (HPR). Within this complex, the APOL1 protein functions as the primary lytic element^4^. TLF provides defense for humans, gorillas, baboons, and select individuals against prevalent African trypanosomes, a condition known as African sleeping sickness, which is caused by *Trypanosoma brucei*^5–7^. Two genetic variants of APOL1 emerged within the human population of sub-Saharan Africa rapidly disseminated across all African populations due to their ability to offer heightened defense against the virulent subspecies of trypanosomes known as *T.brucei rhodesiense*^8, 9^. These genes variants are G1 coding variant which encompasses two non-synonymous single nucleotide polymorphisms (SNPs), while G2 involves an in-frame deletion of the amino acids N388 and Y389 (**Figure 1A**).

**Figure 1.**
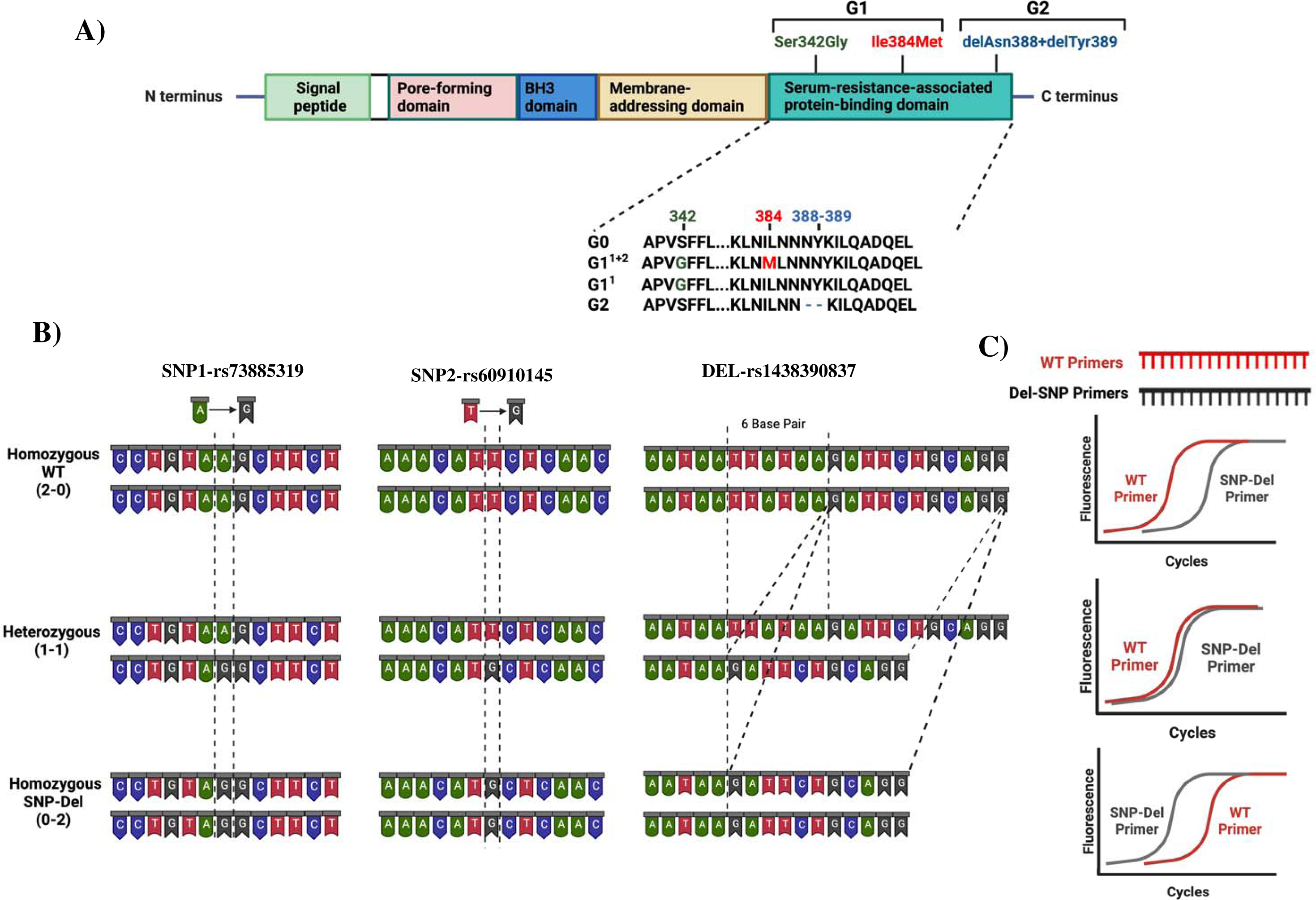
Risk Variants of APOL1 gene. **A)** Graphical representation of the APOL1 protein depicting its domains and location of G1 and G2 variants in the SRA domain **B)** Illustration of base changes at individual SNP and deletion sites. The DNA structures for homozygous wild-type (2-0), heterozygous (1-1), and homozygous SNP/deletion (0-2) variants are displayed across three panels. Locations and genotypes are detailed with dotted lines **C)** Predicted results from qPCR analysis utilizing our probe-independent Real-Time quantitative PCR approach.

While these variants have shown effectiveness in enhanced combating on *T.brucei* infections, they are also linked to a range of kidney-related disorders. These variants increase susceptibility to inflammation, aggravation of lupus glomerulonephritis, focal segmental glomerulosclerosis (FSGS), chronic kidney disease (CKD), end-stage renal disease (ESRD) as well as early rejection of transplanted organs^10–12^. These genetic variants influence the survival of kidney transplant recipients by increasing the risk of allograft dysfunction. Patients carrying one or two risk alleles are more likely to experience kidney dysfunction, potentially compromising transplant outcomes compared to patients without these risk alleles^13^. These variants cause collapsing glomerulopathy, podocytopathy, and tubulopathy via mechanisms for podocyte injury including cationic pore-forming ability, altered autophagy, and direct toxicity^14–16^. The toxicity of risk variants is associated with background haplotype. Based on these 2 SNPs and 1 deletion, all risk alleles can be classified as no-risk allele (G0/G0), one-risk allele (G0/G1, G0/G2) and two risk allele genotypes (G1/G1, G1/G2, G2/G2)^17^. Nevertheless, it is important to consider that offspring inherit one allele from four potential haplotypes from each parent. The prior division of APOL1 genotypes, such as 2 groups for one-risk alleles and 3 groups for two-risk allele, could result in misinterpretation when assessing the impact of APOL1 variants and categorizing APOL1-related kidney diseases. Thus, a more nuanced subgrouping within each risk category might be necessary, based on an analysis of individual SNPs at G1 position and their combinations with deletion variant at G2 (**Figure 1A**).

Genetic testing and probe-reliant PCR screening are currently used for detection of variants, alongside costly and labor-intensive mass spectrometry methods. This underscores the need for quicker, more cost-effective ways to identify variants, especially for screening patients for potential risks in clinical settings. To address this, we have developed a probe-independent quantitative polymerase chain reaction (qPCR) technique for identifying APOL1 variants (**Figure 1B**), primarily aimed at our transplant cohorts, which mainly consist of African-American patients.

## METHOD

### Study Design and Participants

This study included samples from patients who had kidney transplant surgery. Briefly all patients admitted to the Transplant Institute in Methodist University Hospital from May 2021 to May 2023 were considered for inclusion in this prospective study. The study comprised patients who had elective transplant surgery. 171 patients were included in our sample biobank upon given written informed consent for participation. Samples were collected before and after surgery for patients and after receiving the organ from donors. Following hospital discharge, patients were prospectively followed up for 12-months (weeks 1, 2, 3, 4, and months 2, 3, 6, 9 and 12). All biochemical and patient outcomes were recorded.

The study protocol was approved by the local ethics committee by The Institutional Review Board of the University of Tennessee Health Science Center (IRB Approval Number: 20-07838-XP). All clinical investigations were conducted in accordance with the Declaration of Helsinki. Demographic characteristics and laboratory findings were collected.

### DNA Isolation from Patient Bloods

We isolate genomic DNA from PBMC samples of recipients. Blood samples were collected from patients into tubes with K2 EDTA additive before the surgery to get PBMCs with Ficoll gradient method. Blood samples were initially spun at 2200 G for 30 min and separated from plasma. After this step, they were resuspended with PBS and added onto equal amount of Ficoll solution slowly in a 15 mL centrifuge tube to form 2 separate layers and samples were centrifugated at 2200 G for 15 min without brake. After this step, cloudy buffy coat layer was formed and transferred into new centrifuge tube and collected PBMCs were washed with PBS. PBMCs was either frozen with 10% DMSO in FBS solution or taken to DNA isolation, freshly. PBMC samples were resuspended in 250 µL SDS lysis buffer (100mM NaCl, 50 mM Tris-HCl, 5mM EDTA, 1% SDS), vortexed and sonicated to lyse all cells. After lysis of the cells, they were incubated at 95°C for 5 min and at room temperature on bench for 5 min, respectively. We added 3-4 µL proteinase K (10mg/mL) into samples and incubate at 37°C for 1-2 hours. Following to incubation, 250 µL phenol chloroform isoamyalcohol (PCI) were added and mixtures were vortexed vigorously for 30 sec. Samples were centrifugated at 13000 G for 10 mins and at the end of this spin upper aqueous layer were transferred into a new 1.5 mL tube without disturbing bottom organic phase. We added 500 µL 100% EtOH and 25 µL 3M Na-Acetate to mixture and vortexed. Following centrifugation at 14000 G for 15 min and supernatant was discarded. Remaining pellet was washed with 500 µL 70% EtOH, centrifugated at 14,000G for 5 mins and supernatant were discarded. Samples were dried at room temperature till ensuring all ethanol has been removed. Pellet was dissolved in 100 µL double distilled water. Additionally, 1 µL RNase were added to eliminate any RNA contamination.

### Amplification of APOL1 Gene Segment Using Polymerase Chain Reaction

We first used primers for PCR technique to amplify APOL1 gene segment for SRA domain specifically to include the possible regions of 2 SNPs and 1 deletion region (Primers are represented in Supplementary Table 1). Following the isolation of DNA, we performed a PCR reaction to amplify the SRA domain, utilizing a total of 10 ng of DNA, 2ml of primer mixture (final concentration of 1 mM primers mixture), 10 ml of NEBNext® High-Fidelity 2X PCR Master Mix (New England Biolabs, Ipswich, MA, USA), 2ml of DMSO in dH2O (total 20ml/wheel). We confirmed size of PCR product (∼280 bp) using Agarose gel electrophoresis.

### Sanger Sequencing

Sanger sequencing is used to evaluate DNA products with identifying all deoxynucleotide triphosphates individually and allows to investigate specific sequences for SNPs, mutations, deletions, and mismatches. Amplified products were purified and sent to Eurofins Genomics Laboratories. Sequencing results were evaluated with Snap Gene Viewer application for MAC (GSL Biotech LLC, San Diego, CA). We also developed scoring system for each patients using sanger results; i)if a singular peak corresponding to the wild-type (WT) is seen at the SNP region, or if there’s a full 6-bp sequence present without any perturbation at G2 location; we scored as 2-0 ii) then overlapping peaks are noticed at the SNP locations or overlapping nucleotides appear on or following the deletion region, we assign a score of 1-1, iii)if there is a single peak with modified nucleotide at the SNP location, and definitive 6-bp deletion on sanger chromatogram at G2 location; we assign a score of 0-2. These scores were used to evaluate outcomes of transplant patients.

### qPCR to Detect APOL1 Variants

Quantitative real-time PCR has been conducted to find the ΔCt values between WT and SNP/Del primers. Briefly, we construct WT primers and “SNP/Del primers” for two SNP loci and 6-bp deletion locus. PCR products were run 6 sets of reaction to detect SNP/Del and WT gene for each patient (**Figure 1.B-C).** Initially we used “unmodified exact sequence” primers to match WT, SNP or Del sequences. After finding no expression differences, we used “modified primers”.

We used the following formula for qPCR setup for each reaction. 1 ml PCR product, 10 ml of 2x SYBR Green qPCR master mix (PowerTrack SYBR Green Master Mix, ThermoFischer Scientific, Waltham, MA, USA), 2ml of primer mixture (final concentration of 1 mM primers mixture) in dH2O (total 20ml/wheel). All primer sequences are represented in Supplementary Table 1. We checked both scores of Sanger sequencing and qPCR results to confirm that both scores are similar.

### Statistical Analysis

Statistical analysis was carried out using Graph Pad Prism for Mac software. Kidney function results, which exhibited a normal distribution based on the Kolmogorov–Smirnov test, are presented as mean±standard deviation (SD). To compare patient groups with 0, 1, and 2 Risk Alleles, we conducted separate one-way ANOVA tests, followed by post-hoc Tukey tests with Bonferroni correction at each time point. Additionally, we employed repeated measures ANOVA to compare follow-up data within each risk allele category. Statistical significance was determined as P < 0.05.

## RESULTS

### Evolving strategies in APOL1 Variant Classification: A Haplotype-Centric Model

The stratification of patients based on their APOL1 variant status is critical for anticipation of clinical progress of patients in APOL1-related research, particularly in extensive cohort studies like APOLLO ^18^. Current classification systems generally categorize patients into three groups, depending on the number of risk alleles (0-risk, 1-risk, 2-risk), or into six groups, when the genotypes of individual alleles are considered (G0/G0 for 0-risk, G0/G1 and G0/G2 for 1-risk, and G1/G2, G1/G1, and G2/G2 for 2-risk alleles). Here, we introduce a novel classification approach that considers the haplotypes inherited from each parent and potential combinations. In this new framework, we employ a scoring system that focuses on two specific SNP locations and deletion locus. This offers a more nuanced way of classifying APOL1 variant status, which could lead robust studies 1and possible estimation of patient progression after transplantation. Considering the equal inheritance of genetic material from each parent, we posit the possibility of attributing a scoring mechanism to individual SNPs and deletions. In this proposed model, a score of 2-0 is designated when both DNA sequences present the WT allele. In contrast, a score of 1-1 is given when one DNA sequence displays the WT allele and the other has a SNP or deletion. Lastly, a score of 0-2 is assigned when both DNA sequences stand a SNP or deletion. We performed this scoring for each SNP and deletion so that we get 6-digit code for each patient. For instance, a score of 20-20-20 would represent a WT status for both SNPs and deletion, corresponding to the genotype G0/G0. With this new coding system, we obtained a total of 10 different groups (1 for 0-risk, 3 for 1-risk, and 6 for 2-risk allele, **Figure 2**).

**Figure 2.**
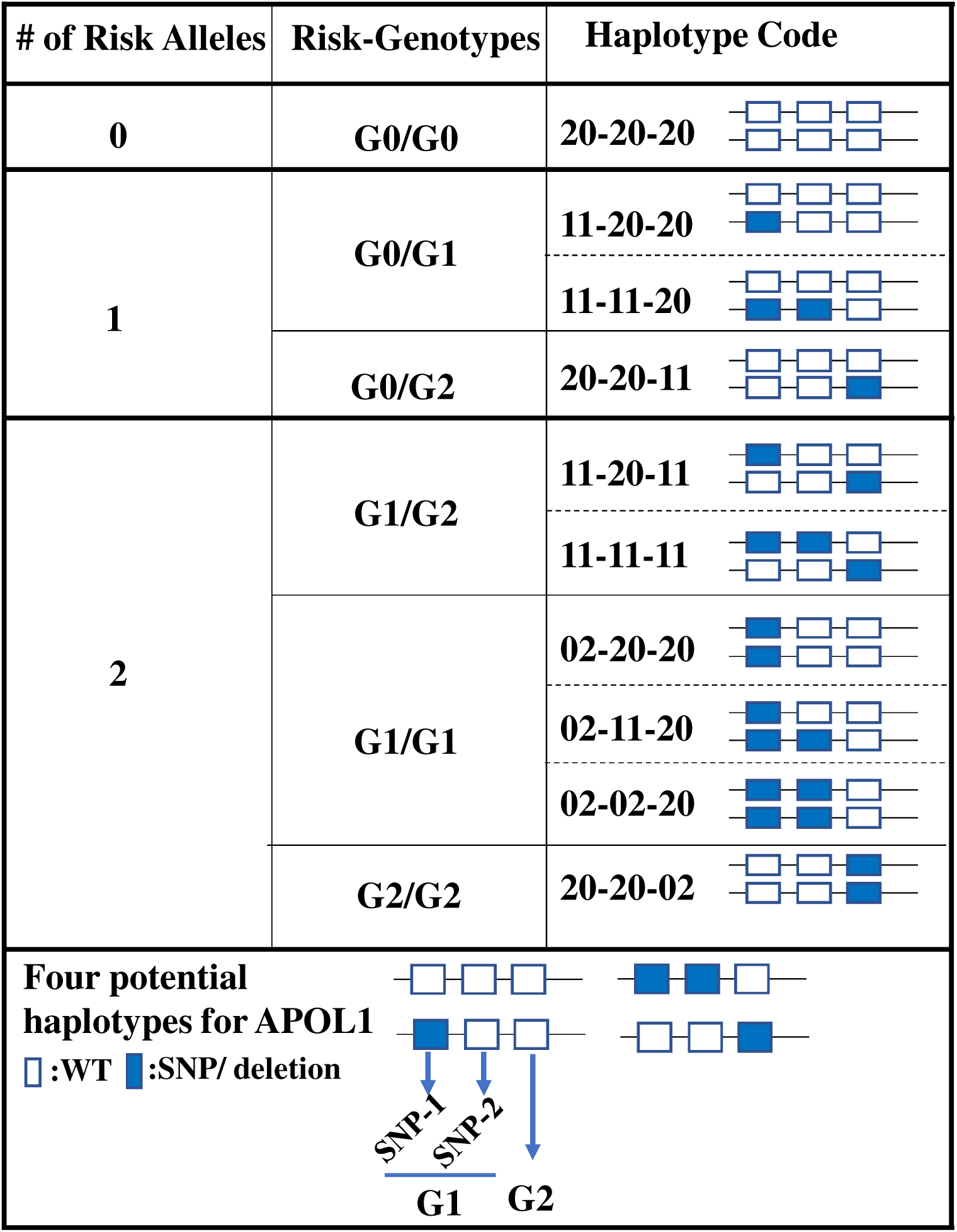
Haplotype-centric classification approach for APOL1 variant status. This figure illustrates the proposed classification model for patients based on their APOL1 variant status. Three primary risk categories (0-risk, 1-risk, 2-risk) are further divided into a total of 10 groups using a novel 6-digit scoring system focusing on APOL1 haplotypes.

### Demographic Characterization of Donors and Stratification of APOL1 Genetic Variants Through Sanger Sequencing

In our kidney transplant cohort, we obtained 155 donor (16 paired, 139 individual kidney transplant) organs and 171 recipients. Within the donor group, 112 individuals identified as Caucasian, 33 as African-American, and 10 belonged to other racial backgrounds. On the other hand, the recipient cohort included 143 African-American, 24 Caucasian, and 4 individuals from other racial backgrounds (**Figure 3.A-B**, **Table1**). Given the higher proportion of African-American patients in our recipient cohort, we anticipated a greater prevalence of APOL1 risk variants. With analysis of distinct peaks observed in the Sanger chromatograms, we successfully ascertained the genotype at each respective position for SNPs and deletions. (**Supplementary Figure 1**). Sanger sequencing results revealed that 82 recipients (48%) were devoid of any risk alleles (G0/G0), while 58 recipients (34%) had one risk allele and 31 recipients (18%) had two risk alleles.

**Figure 3.**
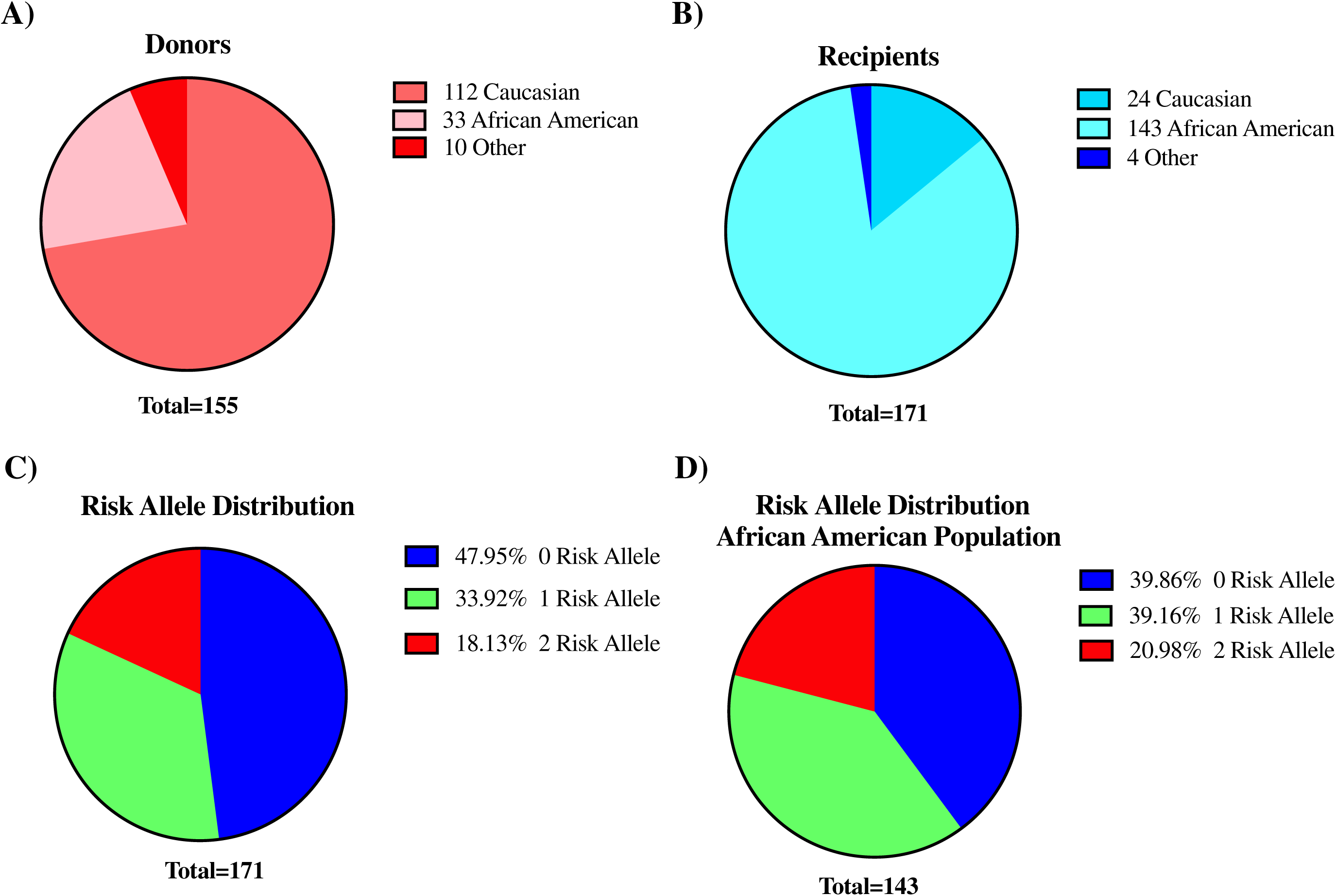
Profile of donor and recipient in our cohort and overview of risk metrics for recipients. Distribution of donors **A**) and recipients **B**) based on their ethnicity. **C)** Distribution of APOL1 risk variants among the 171 recipients in our study **D**). Risk Allele distribution for only African American recipients (n:143) in the cohort. Blue:0-risk, green:1-risk, red:2-risk.

**Table.1:**
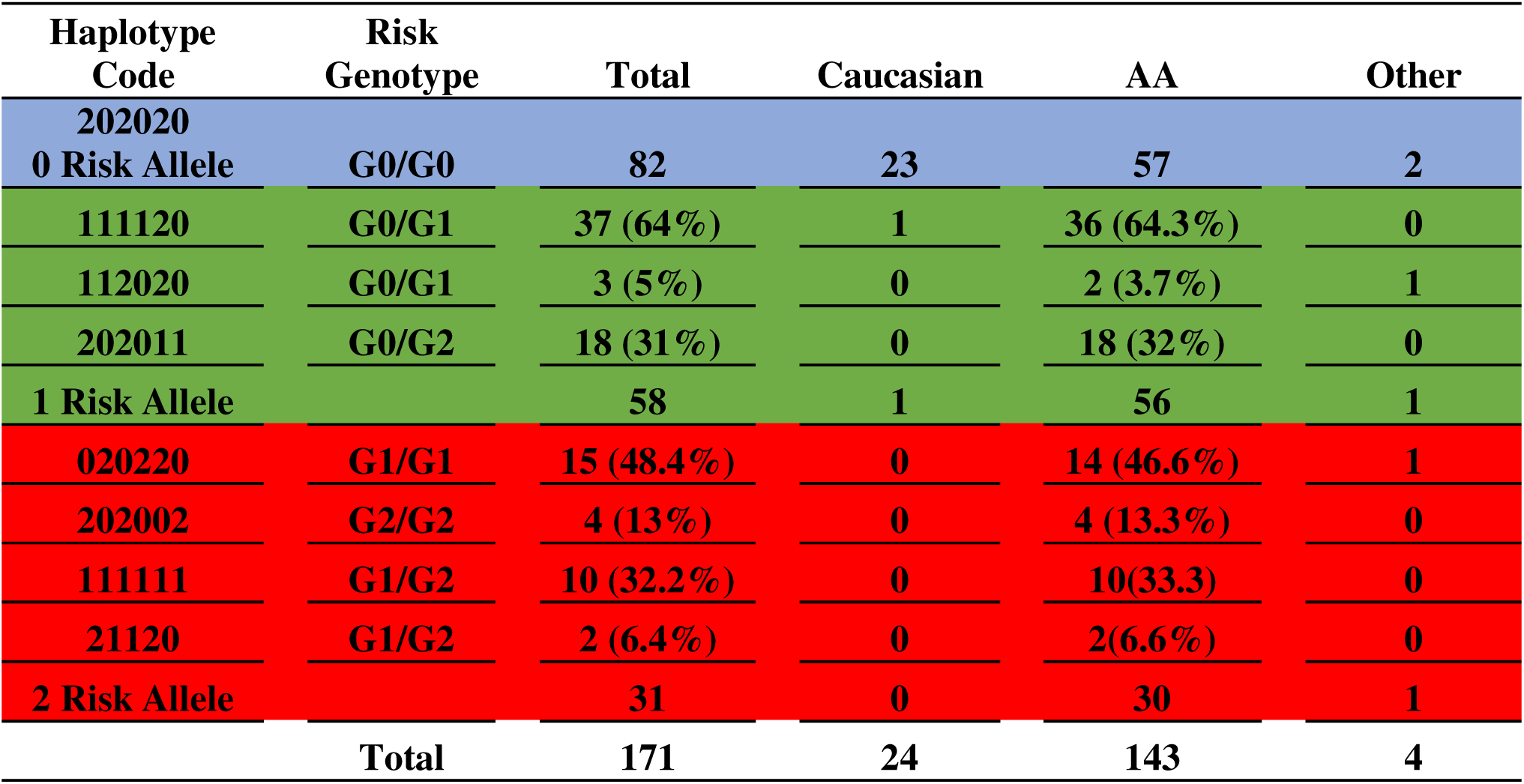
APOL1 Haplotype Codes and Ethnicities of Patients.

We further stratified our patients based on both their APOL1 risk genotype and APOL1 haplotype codes. In this cohort, we observed the presence of all three conceivable haplotype codes associated with 1-risk allele, with haplotype 11-11-20 manifesting at the highest frequency (64%). Similarly, among recipients harboring dual risk alleles, only four of the six putative haplotypes were detected. The haplotype 02-02-20 emerged as the most prevalence (48.4%). It is noteworthy that the haplotypes 02-20-20 and 11-20-11 were absent in our dataset due to the limited number of patients in the cohort. In our recipient cohort, one individual of Caucasian descent and 1 individual of another racial background each carried one-risk allele, while another individual from different racial patient carried two risk allele. Within the African-American subset of cohort, 57 (40%) of recipients had no risk alleles, 56 (39%) had one risk allele, and 30 (21%) had two-risk alleles (**Figure 3C-D and Table 1**).

### Probe-Independent Quantitative PCR as a Robust Alternative to Genetic Testing and Sanger Sequencing for Variant Identification

#### SNP1(rs73885319) region and SNP-2 (rs60910145) region

Following the initial identification of APOL1 variants through clinical genetic testing and Sanger sequencing, we selected a subset of patients with pre-identified genotypes to serve as control. Our initial approach involved designing allele-specific primers to distinguish the WT allele from the SNP allele. This design was predicated on the concept that a single nucleotide difference at the 3’ end of primer oligos could significantly alter the PCR efficiency. Our objective was to detect this efficiency difference in real-time quantitative PCR as a ΔCt value, thereby eliminating the need for an internal probe (**Figure 1C**). We termed this set of primers as "unmodified primers," which differed only by a single base at the SNP locus. Specifically, for the SNP1(rs73885319) region, we implemented a single base change at the end of the sequences (A→G), and for the SNP-2 (rs60910145) region, we used a single base change at the end of the sequences (T→G). Initial qPCR assays using these primers yielded similar expression levels for both WT and SNP genotypes, as illustrated in **Figure 4 A-B** (left part in each section). It became evident that a single nucleotide changes at the primers’ 3’end was insufficient for distinguishing between the groups using qPCR. Subsequently, we opted to modify the primers by introducing a second base pair alteration adjacent to the SNP location to affect the binding efficiency of the primers to their target regions. In this modified format, we switched the second base from a purine to a pyrimidine pair for SNP1 or keep it as pyrimidine but change to cytosine for SNP2, thereby enhancing our ability to detect differences between the WT and SNP genotypes. So, we changed the last two nucleotides from AA to TA for WT primer and AG to TG for SNP primer for SNP1 region. Similarly, we changed the last two nucleotides from TT to CT for WT primer and TG to CG for SNP primer for SNP2 region. We called these primer pairs as “modified primers” (**Figure 4A-B, Supplementary Figure 2**).

**Figure 4.**
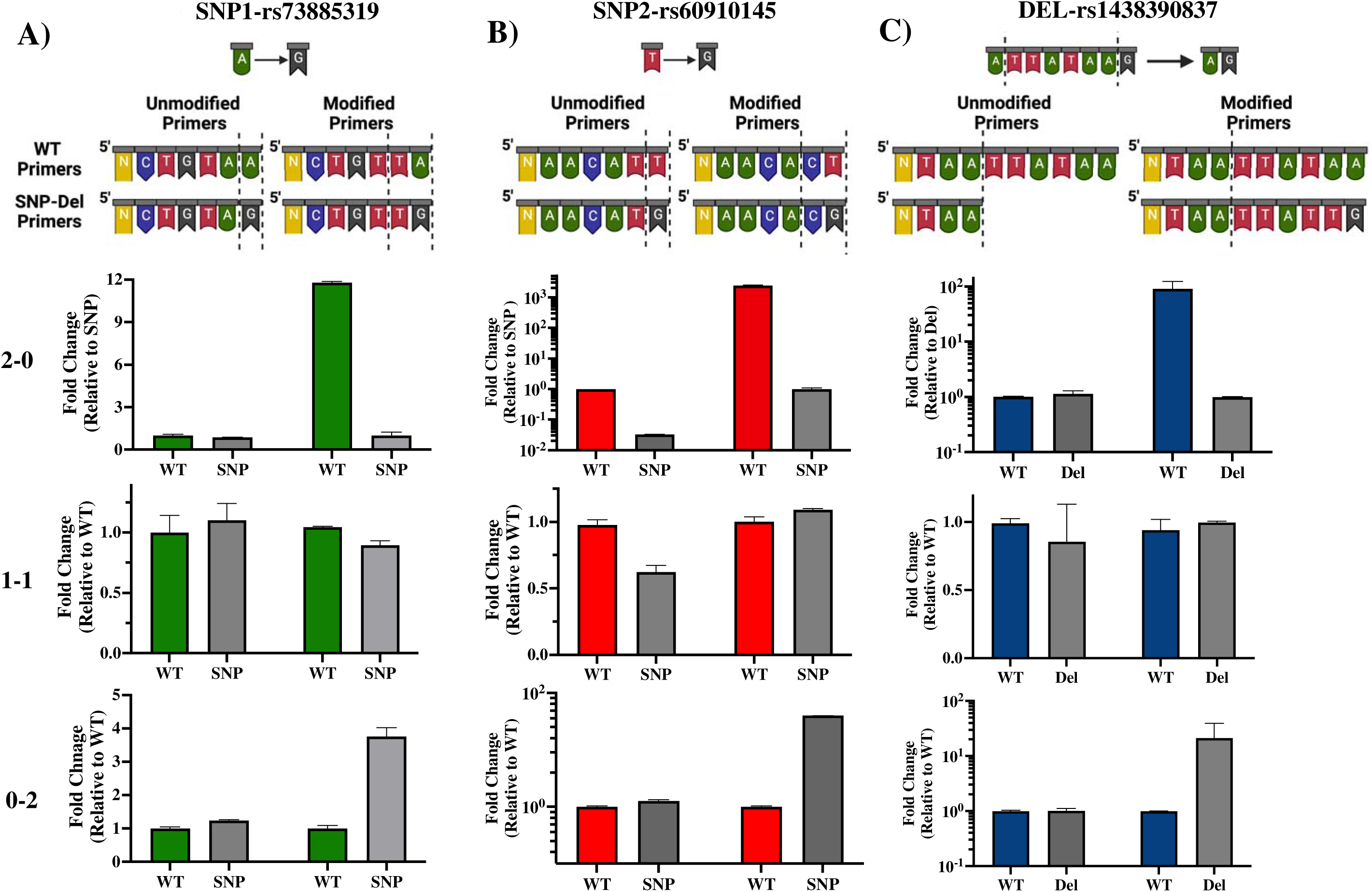
Development and validation of allele-specific qPCR primers for APOL1 variant detection. Top panel: Illustration of standard “unmodified” or custom-designed “modified” primer sequences for SNP-1 (A), SNP-2 (B) and deletion (C) locus. Altered or deleted base pairs demarcated by dotted lines. **Bottom panel**) Quantitative analysis of qPCR data illustrates the homozygosity or heterozygosity variants. 2-0: homozygote WT, 1-1: heterozygote, 0-2: homozygote SNP/deletion (N: Remaining bases in the primer sequence)

In this new qPCR setting, we observed a minimum 4-fold differential expression (ΔCt>=2) between the WT primers and SNP primers when using homozygous WT or homozygous SNP genomic DNA for both SNP loci. In qPCR where we used heterozygous DNA, the differences in expression levels between the two primers were uniform and did not exceed a 4-fold difference (**Figure 4 A-B**, right side in each panel, **Figure 5A-B, Supplementary Figure 2**). Upon optimizing our primers for specific SNP detection, we proceeded to evaluate all recipient genotypes to ascertain the ΔCt values, which were then utilized to calculate the threshold value. As shown in **Figure 5 A-B**, a distinct separation in ΔCt values were evident among all three genotype groups.

**Figure 5.**
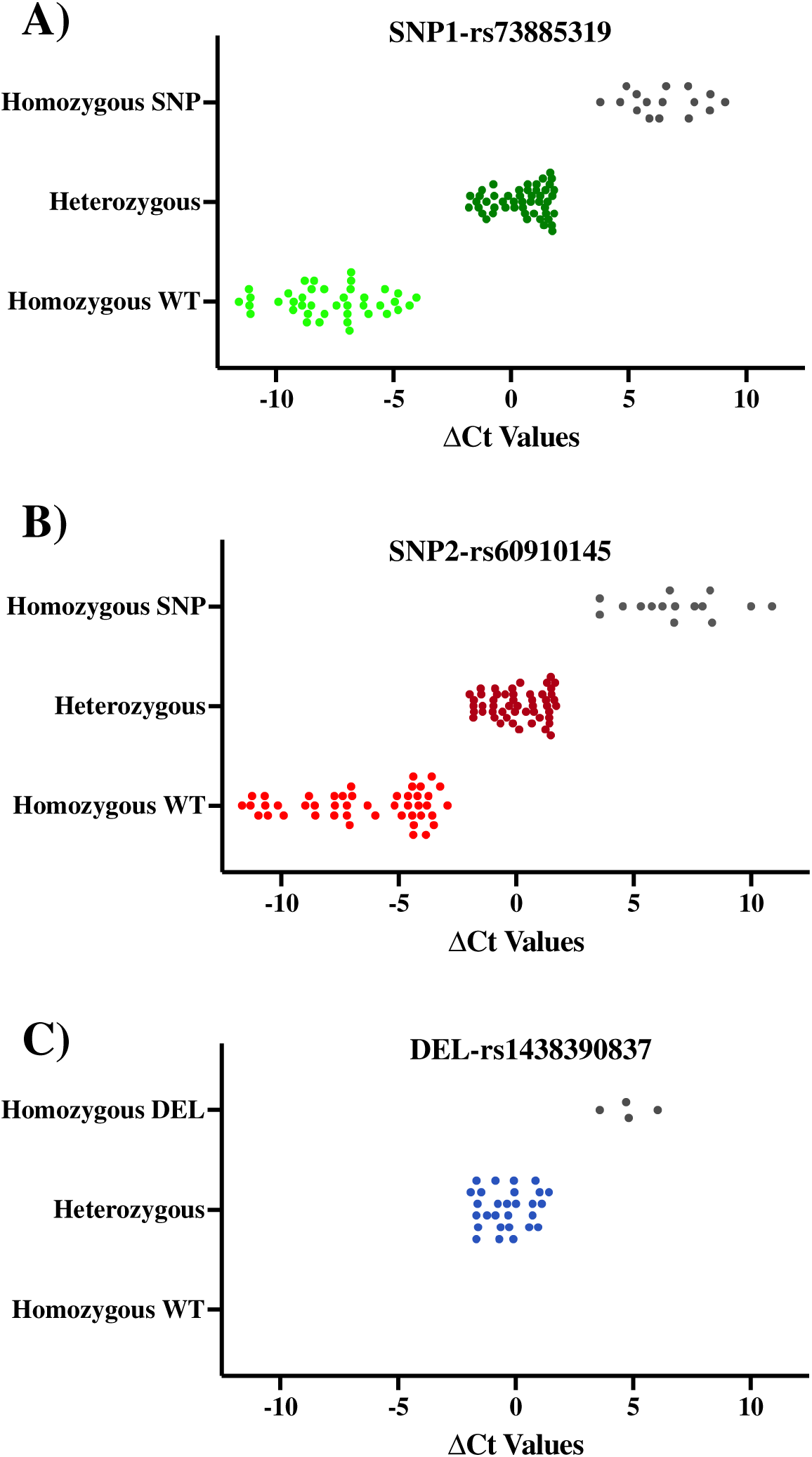
ΔCt Values of QPCR Results for SNPs and Deletion. qPCR results of each SNP and deletion which are classified based on Sanger Sequencing results. Each data point shows mean of ΔCts values (ΔCt=Ct value with modified WT primer-Ct value with modified SNPs/Del primer). Lighter color: Homozygous WT, Darker color: Heterozygous, Gray: Homozygous SNP or Deletion

### Deletion rs143830837 Region

Our PCR reaction was initially configured using a common forward primer, along with reverse primers with exact match designed to detect the presence or absence of 6-bp deletion identified as “TTATAA” (these primers are termed “Unmodified Primers”). In this quantitative PCR setup, we did not observe any notable expression differences between patients with WT DNA and those with 6-bp deletions in their DNA (**Figure 4C**, left part in each section, **Supplementary Figure 2**). Drawing on our experience with SNP primers, where we distinguished between wild-type and SNP DNA by modifying 2 base pairs at the 3’ end of the primer, we applied the same approach to the deletion region. We chose to extend the primers to create a 2-base pair mismatch between them. However, within the rs143830837 Region, there is a repetitive sequence of adenine and thymine bases (5’-ATAA). We opted to extend our WT primers by incorporating the exact sequence from the 3’ end. For the Del primers, we used the same template as the WT primer and extended it with the exact sequences of DNA containing the deletion, resulting in 2 base differences (ending with “AA” in the WT primer and “TG” in the Del primer). These modified primers were referred to as “Modified Primers”.

In this qPCR setup, we observed expression differences of at least 4-fold for the WT genotype compared to the deletion genotype in the 0-2 and 2-0 genotyping. In the 1-1 genotyping, the differences in expression levels between both primers were equal and less than 4-fold (**Figure 4-C**, right part in each section, **Figure 5C, Supplementary Figure 2**). Original quantitative PCR graphs for SNP loci and deletion locus are represented in **Supplementary Figure 3**.

### APOL1 Variants Show No Early Impact on Renal Function Metrics in Kidney Transplant Recipients

As patient evaluation before and after surgery at day-1 and -2, as well as week-1, and months 1, 3, 6, 9 and 12 to monitor their recovery progress, the levels of Blood Urea Nitrogen (BUN), creatinine, urinary protein, and Estimated Glomerular Filtration Rate (eGFR) were measured to assess the functionality of the transplanted kidneys. We categorized the results based on the presence or absence of risk alleles. Over the course of the first month, BUN, creatinine, and urine protein levels showed a significant decrease before stabilizing at 3-, 6-, 9- and 12-month follow-ups (p<0.05). Similarly, eGFR levels showed a progressive improvement during the first month, which then remained stable during the 3-, 6-, 9- and 12-month follow-ups (p<0.05). Kidney functions were similar and were improving for all risk allele groups and were steady after 3 month follow-ups. Importantly, no notable differences were observed in these renal functional indicators when stratifying the recipients based on APOL1 risk allele status (**Figure 6**, **Supplementary Figure 4**).

**Figure 6.**
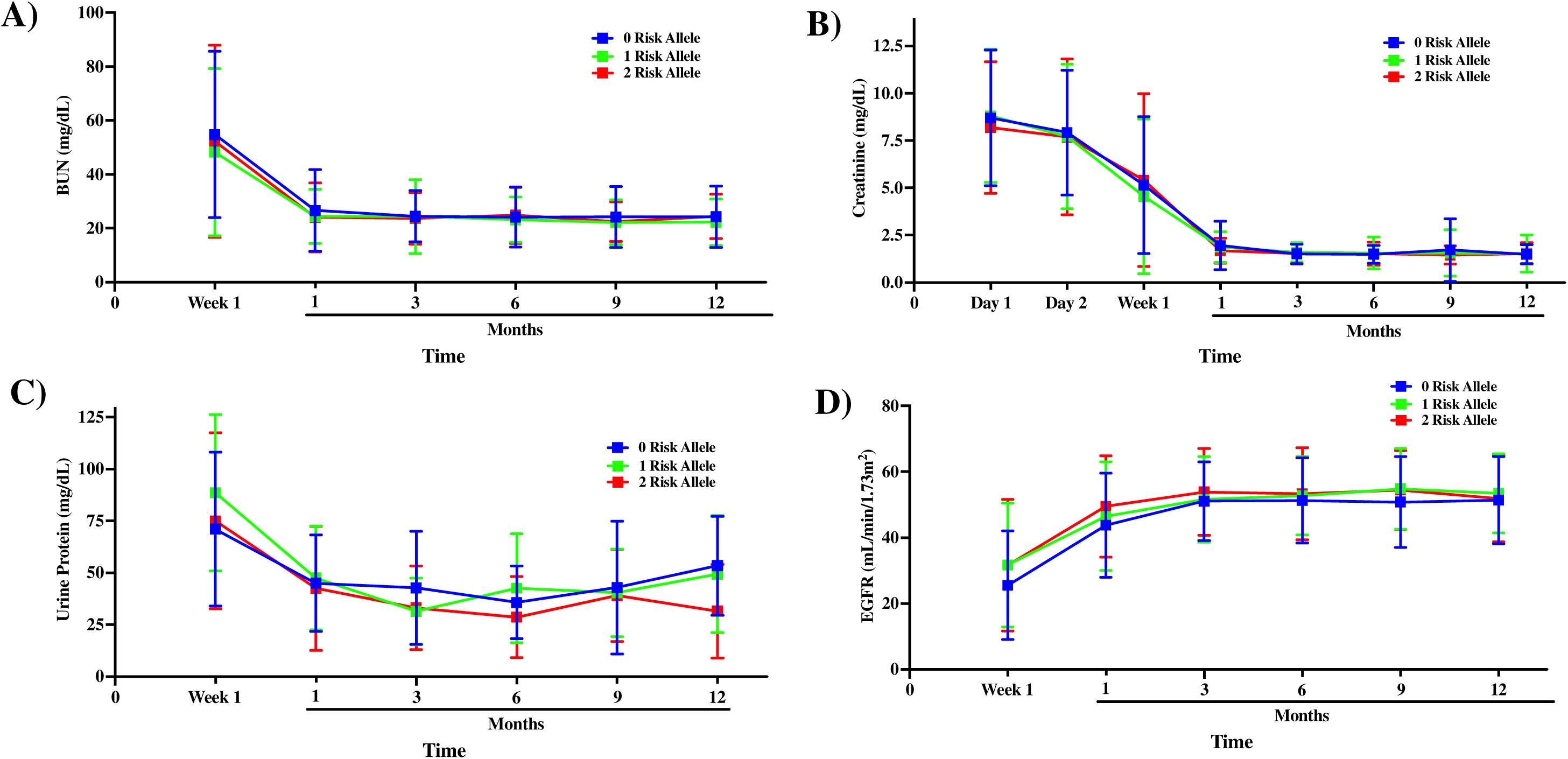
Comparison of Kidney Function Among APOL1 Risk Alleles Post Transplantation. Kidney functions were compared after transplantation to up to-1-year follow-ups; **A)** blood BUN**, B)** Creatinine, **C)** Urine protein, **D)** eGFR progression Blue:0-risk, green:1- risk, red:2-risk.

## DISCUSSION

This study delves into APOL1 variants, their identification, patient genotype classification, haplotype coding, and implications for kidney transplant recipients. We introduce a novel haplotype-centric model for APOL1 variant classification and employ quantitative PCR as an alternative genotyping method. Furthermore, we analyze early post-operative renal function metrics in kidney transplant recipients based on their APOL1 risk allele status. These findings offer valuable insights into the expanding field of APOL1 and kidney disease research.

Our innovative haplotype-centric model marks a significant advancement over traditional APOL1 classification methods, which were limited to a few risk allele categories or genotypes. Our approach introduces a unique 6-digit code, enabling a wide range of haplotype combinations for a more nuanced understanding of APOL1-related risks. Within our patient group, we successfully identified 8 out of 10 possible haplotype combinations, with the remaining two eluding us, likely due to our cohort’s limited size.

Currently various methods are employed to identify APOL1 risk variants in individuals, including the measurement of APOL1 variants bound to HDL particles. APOL1, a structural protein of HDL, is cleared from the bloodstream by the kidneys. Assessing mutant APOL1-HDL proteins’ quantity and quality may aid in disease estimation. In a recent study involving 3450 individuals, liquid chromatography-mass spectrometry (LC-MS) was used to measure plasma APOL1 variant levels, but no association with kidney function was found. The study concluded that circulating APOL1 levels may not correlate with mutant APOL1-HDL protein forms^19^. Additionally, another group identified APOL1 gene variants with blood and serum measurements using LC-MS, analyzing surrogate peptides to identify different gene variants^20^.

The probe-based method is another way to identify APOL1 gene variants. It involves isolating DNA from blood samples and using fluorescently labeled detection probes along with PCR and a 5’-nuclease assay. When a probe matches the target DNA containing APOL1 variants, it’s degraded by 5’-nuclease polymerase activity, releasing the reporter dye and producing a detectable fluorescent signal. The TaqMan SNP Genotyping Assay, commonly used in clinical settings, is a standard choice for this method^21^.

Our study utilized custom-designed primers and quantitative PCR for APOL1 gene variant identification, offering quicker and cost-effective results. We extracted DNA from whole blood, then used variant-specific primers to quantify threshold cycle (Ct) values. Validation through Sanger sequencing yielded a 100% success rate, making this method accessible for standard research laboratories.

Identifying APOL1 gene variants in potential kidney donors and transplant recipients holds crucial health implications. For donors, possessing these variants increases the risk of kidney dysfunction and end-stage renal disease, potentially making kidney donation a risky endeavor. Moreover, it can impact recipients, leading to early allograft dysfunction and rejection. A study by Reeves-Daniel et al. revealed significantly lower graft survival rates in donors with 2 APOL1 gene variants compared to those with 1 variant (50% vs. 75% over 3.5 years). Notably, having 2 APOL1 variants poses a higher risk (HR-3.84) than HLA mismatch and cold-ischemia time (HR-1.52 and HR-1.06, respectively)^22^.

When considering the genotype of transplant patients, in a 5-year retrospective study including 119 African-American transplant recipients, those with 2 APOL1 gene variants showed a similar allograft survival rate to those with 1 variant (around 50% survival rate for both groups), regardless of donor genotype. Despite their heightened risk of native kidney disease, allograft outcomes remained comparable^23^. In a recent research by Zhang et al. presents conflicting results. In the Genomics of Chronic Allograft Rejection (GOCAR) study, patients with 0 APOL1 risk alleles had higher transplant survival rates than those with 1 allele, and 1 allele carriers had better survival than those with 2 alleles during 7-year post-transplantation follow-ups. The Clinical Trials in Organ Transplantation (CTOT) study showed a similar pattern in survival curves, though differences weren’t statistically significant within the groups during 5- year follow-ups^13^.

Our study, which included a substantial sample size of 171 patients primarily of African American descent, revealed a high percentage of APOL1 variants (18% with 2 Risk Alleles and 34% with 1 Risk Allele) among transplant patients. However, the limitation lies in our short post-transplant follow-up timeframe, necessitating extended data collection for more conclusive results. Based on one-year post-transplant outcomes, recipient APOL1 status doesn’t seem to significantly impact kidney functions. While our study couldn’t compare all haplotype risk codes as intended, our probe-independent qPCR-based APOL1 genotyping method could facilitate larger multicenter studies. This new haplotype-centric classification could serve as an alternative categorization for APOLLO study^18^ .

### Achievements

Our probe-independent qPCR method effectively detects APOL1 gene variants. It enables us to extract DNA from minimal blood samples and conduct APOL1 genotyping using qPCR within a total timeframe of 3-4 hours. This breakthrough presents a fresh opportunity in both clinical and basic research for identifying APOL1 gene variants.

## Acknowledgement

Individuals have been supported by the institutional start up at Transplant Research Institute (AB, Canan K and Cem K). National Institute of Diabetes and Digestive and Kidney Diseases of the National Institutes of Health (NIH) under award number R01DK117183 and DK132230 (AB) partially support current study.

Author Contributions: Canan K, Cem K and M.D. designed the study. M.D. together with C.W., G.M.R., H.I., and N.M. performed experiment and data analysis. J.D.E., N.N, C.E., H.R., and J.V. helped us to collect biospecimens. Cem K and M.D. wrote the manuscript with feedback of Canan K, M.T. and A.B.

Conflict of Interest Statement: The authors declare no conflict of interest.

## REFERENCES

1. Smith EE, Malik HS. The apolipoprotein L family of programmed cell death and immunity genes rapidly evolved in primates at discrete sites of host-pathogen interactions. Genome Res 2009; 19: 850–858.

2. Monajemi H, Fontijn RD, Pannekoek H, et al. The apolipoprotein L gene cluster has emerged recently in evolution and is expressed in human vascular tissue. Genomics 2002; 79: 539–546.

3. Madhavan SM, O’Toole JF, Konieczkowski M, et al. APOL1 localization in normal kidney and nondiabetic kidney disease. J Am Soc Nephrol 2011; 22: 2119–2128.

4. Raper J, Fung R, Ghiso J, et al. Characterization of a novel trypanosome lytic factor from human serum. Infect Immun 1999; 67: 1910–1916.

5. Molina-Portela MP, Samanovic M, Raper J. Distinct roles of apolipoprotein components within the trypanosome lytic factor complex revealed in a novel transgenic mouse model. J Exp Med 2008; 205: 1721–1728.

6. Pays E, Vanhollebeke B. Human innate immunity against African trypanosomes. Curr Opin Immunol 2009; 21: 493–498.

7. Vanhamme L, Paturiaux-Hanocq F, Poelvoorde P, et al. Apolipoprotein L-I is the trypanosome lytic factor of human serum. Nature 2003; 422: 83–87.

8. Genovese G, Friedman DJ, Ross MD, et al. Association of trypanolytic ApoL1 variants with kidney disease in African Americans. Science 2010; 329: 841–845.

9. Friedman DJ, Pollak MR. Genetics of kidney failure and the evolving story of APOL1. J Clin Invest 2011; 121: 3367–3374.

10. Shetty AA, Tawhari I, Safar-Boueri L, et al. COVID-19-Associated Glomerular Disease. J Am Soc Nephrol 2021; 32: 33–40.

11. Larsen CP, Bourne TD, Wilson JD, et al. Collapsing Glomerulopathy in a Patient With COVID-19. Kidney Int Rep 2020; 5: 935–939.

12. Friedman DJ, Pollak MR. APOL1 Nephropathy: From Genetics to Clinical Applications. Clin J Am Soc Nephrol 2021; 16: 294–303.

13. Zhang Z, Sun Z, Fu J, et al. Recipient APOL1 risk alleles associate with death-censored renal allograft survival and rejection episodes. J Clin Invest 2021; 131.

14. Beckerman P, Bi-Karchin J, Park AS, et al. Transgenic expression of human APOL1 risk variants in podocytes induces kidney disease in mice. Nat Med 2017; 23: 429–438.

15. Olabisi OA, Zhang JY, VerPlank L, et al. APOL1 kidney disease risk variants cause cytotoxicity by depleting cellular potassium and inducing stress-activated protein kinases. Proc Natl Acad Sci U S A 2016; 113: 830–837.

16. O’Toole JF, Schilling W, Kunze D, et al. ApoL1 Overexpression Drives Variant-Independent Cytotoxicity. J Am Soc Nephrol 2018; 29: 869–879.

17. Lannon H, Shah SS, Dias L, et al. Apolipoprotein L1 (APOL1) risk variant toxicity depends on the haplotype background. Kidney Int 2019; 96: 1303–1307.

18. Freedman BI, Moxey-Mims M. The APOL1 Long-Term Kidney Transplantation Outcomes Network-APOLLO. Clin J Am Soc Nephrol 2018; 13: 940–942.

19. Kozlitina J, Zhou H, Brown PN, et al. Plasma Levels of Risk-Variant APOL1 Do Not Associate with Renal Disease in a Population-Based Cohort. J Am Soc Nephrol 2016; 27: 3204–3219.

20. Labcorp: Apolipoprotein L1 Risk Variants. In, 2023

21. Laboratories MC: TEST ID : APOL1. In (vol 2023), 2021

22. Reeves-Daniel AM, DePalma JA, Bleyer AJ, et al. The APOL1 gene and allograft survival after kidney transplantation. Am J Transplant 2011; 11: 1025–1030.

23. Lee BT, Kumar V, Williams TA, et al. The APOL1 genotype of African American kidney transplant recipients does not impact 5-year allograft survival. Am J Transplant 2012; 12: 1924–1928.

