## Supplementary for "Unveiling APOL1 Haplotypes: A Novel Classification Through Probe-Independent Quantitative Real-Time PCR"

**Supplementary Figure 1**

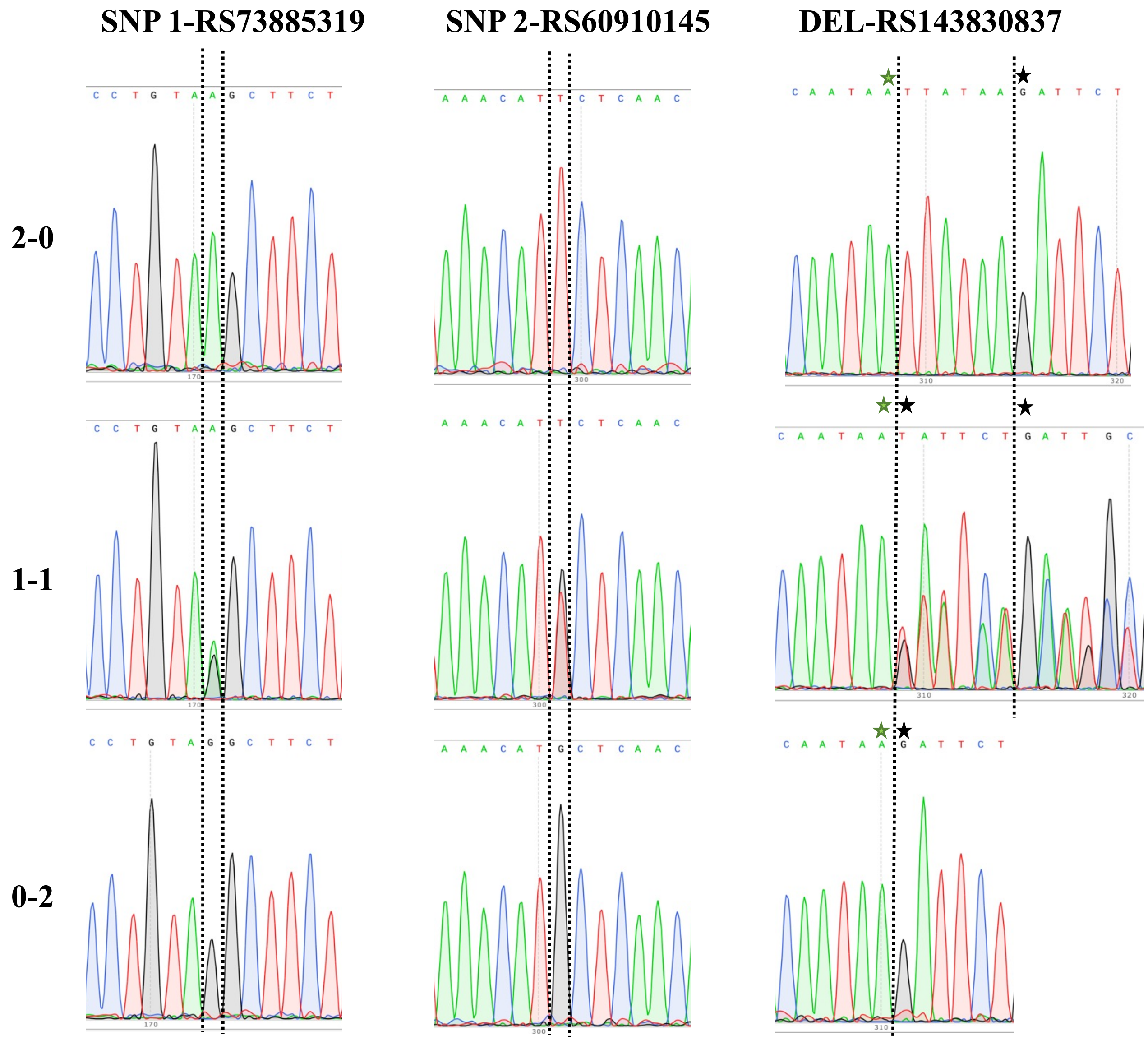


**Supplementary Figure 1** All possible genotypes were determined with Sanger Sequencing for each SNPs and deletion.

#### Supplementary Figure 2

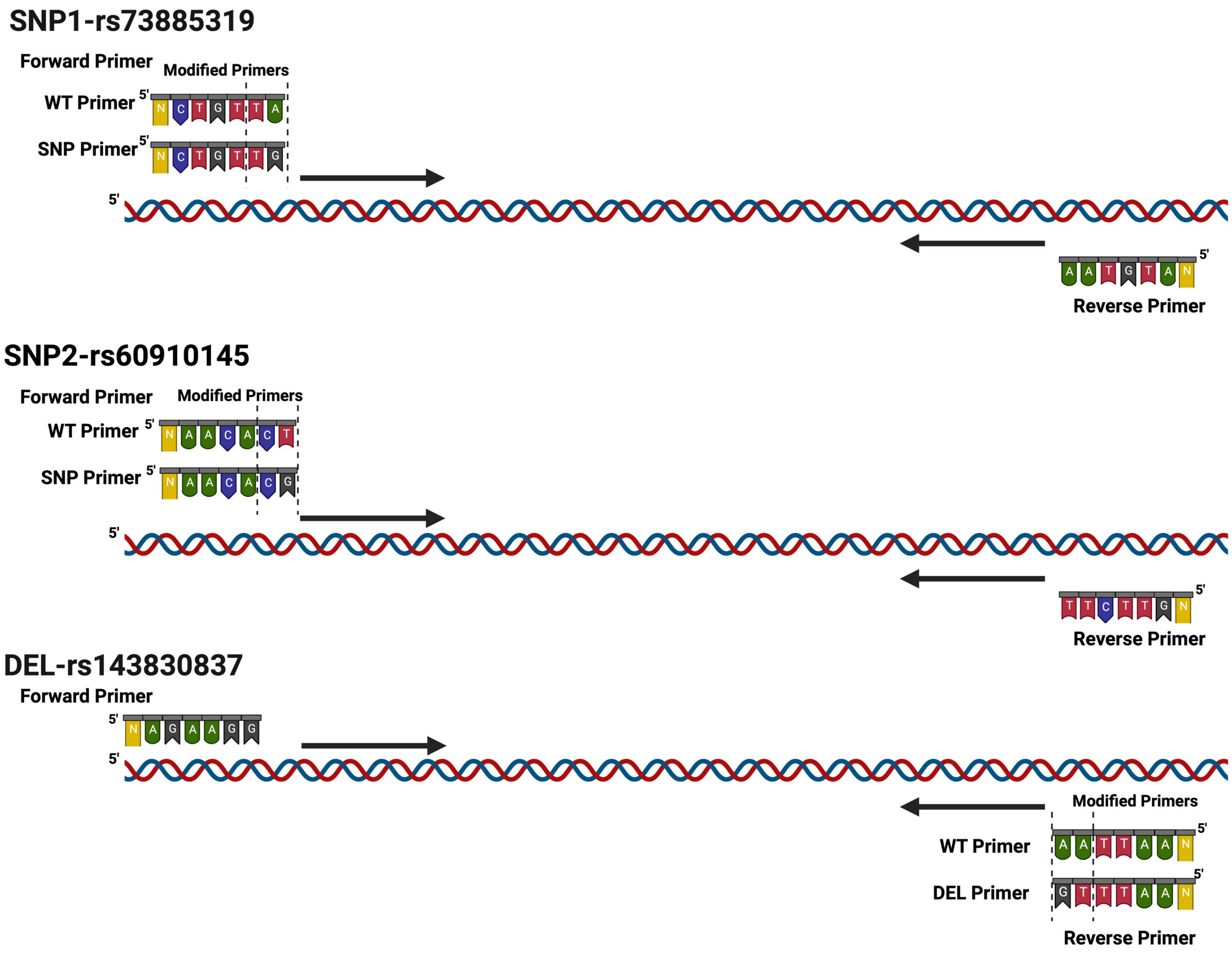


**Supplementary Figure 2** Our primers design for detection of APOL1 gene variants (N: Remaining bases in the primer sequence)

### Supplementary Figure 3

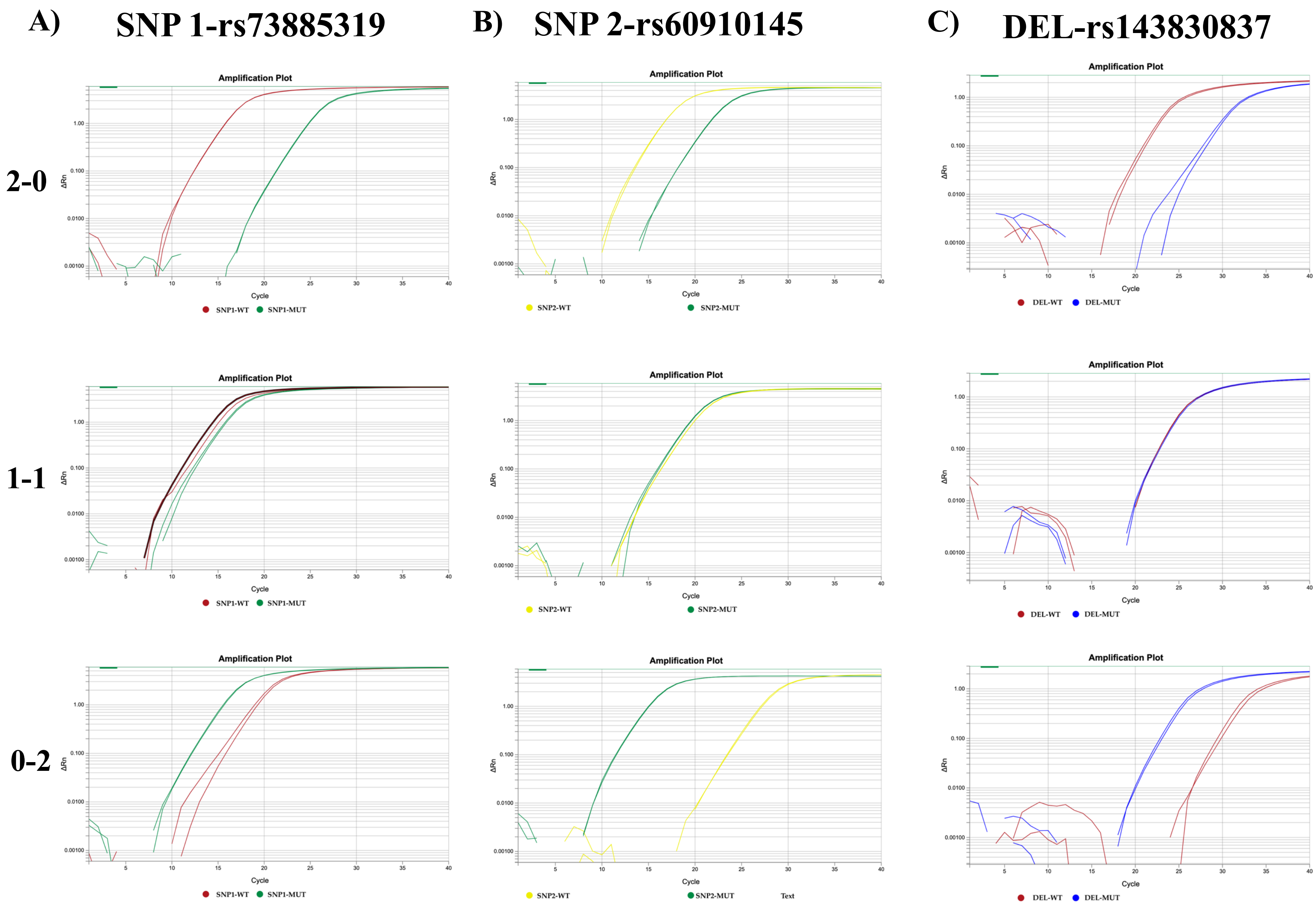


**Supplementary Figure 3** Raw Ct graphs were shown to confirm that modified primers can distinguish all possible genotypes.

### Supplementary Figure 4

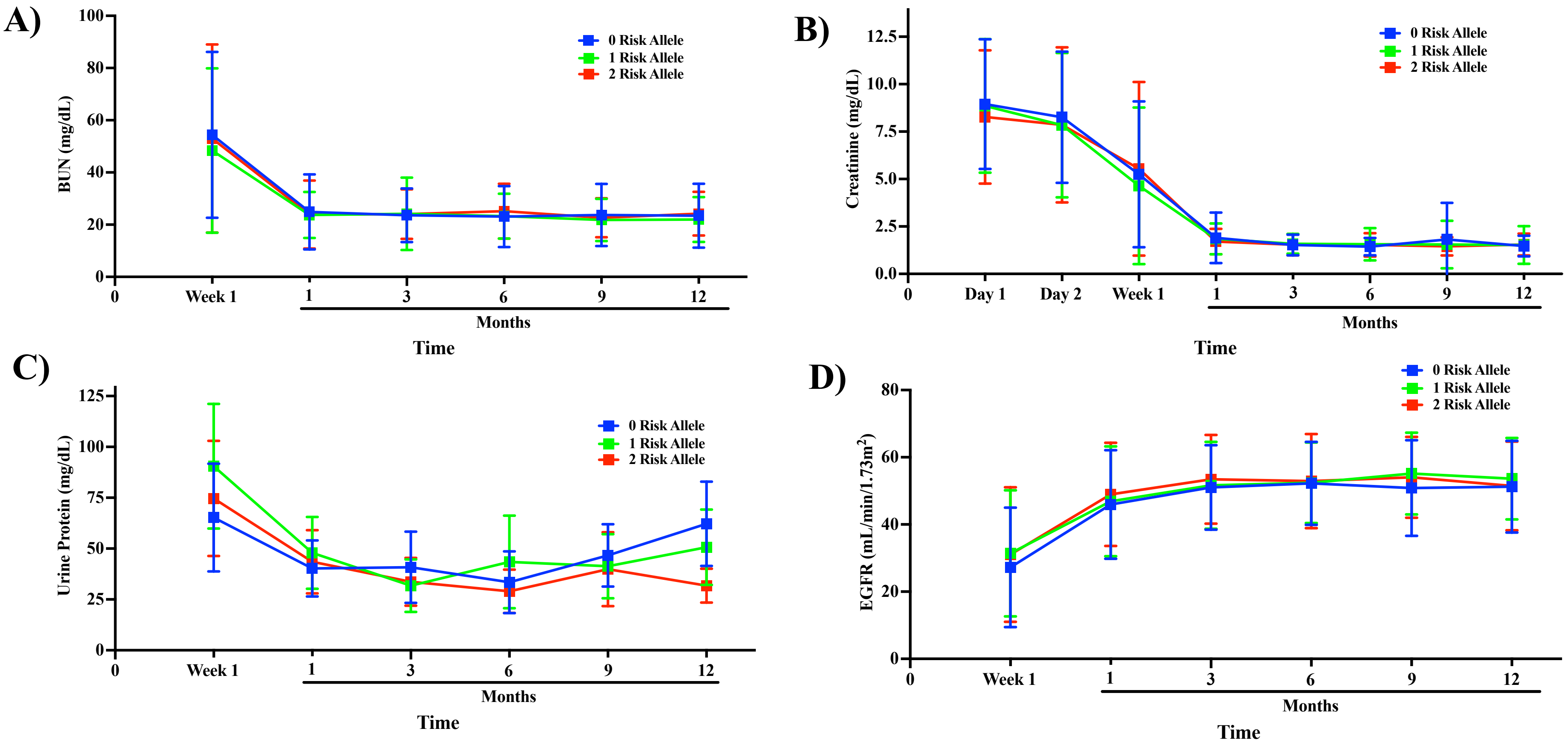


**Supplementary Figure 4** Kidney function was studied in African American recipients based on risk allele carriage. There were no significant differences in kidney function among the groups.
